# Novel pathogen introduction rapidly alters evolved movement strategies, restructuring animal societies

**DOI:** 10.1101/2022.03.09.483239

**Authors:** Pratik Rajan Gupte, Gregory F. Albery, Jakob Gismann, Amy R. Sweeny, Franz J. Weissing

## Abstract

Animal sociality emerges from individual decisions on how to balance the costs and benefits of being sociable. Movement strategies incorporating social information — the presence and status of neighbours — can modulate spatial associations, helping animals avoid infection while benefiting from indirect information about their environment. When a novel pathogen is introduced into a population, it should increase the costs of sociality, selecting against gregariousness. Yet current thinking about novel pathogen introductions into wildlife neglects hosts’ potential evolutionary responses. We built an individual-based model that captures essential features of the repeated introduction, and subsequent transmission of an infectious pathogen among social hosts. Examining movements in a foraging context, widely shared by many species, we show how introducing a novel pathogen to a population provokes a rapid evolutionary transition to a dynamic social distancing movement strategy. This evolutionary shift triggers a disease-dominated ecological cascade of increased individual movement, decreased resource harvesting, and fewer social encounters. Pathogen-risk adapted individuals form less clustered social networks than their pathogen-risk naive ancestors, which reduces the spread of disease. The mix of post-introduction social movement strategies is influenced by the usefulness of social information and disease cost. Our work demonstrates that evolutionary adaptation to pathogen introductions and re-introductions can be very rapid, comparable to ecological timescales. Our general modelling framework shows why evolutionary dynamics should be considered in movement-disease models, and offers initial predictions for the eco-evolutionary consequences of wildlife pathogen spillover scenarios.

## Introduction

Animal sociality emerges from individual decisions that balance the benefits of associations against the costs of proximity or interactions with neighbours (Tanner and Jackson 2012; Gil et al. 2018; Webber and Vander Wal 2018; Webber et al. 2022). While such associations can inadvertently or deliberately yield useful social information about resource availability (Danchin et al. 2004; Dall et al. 2005; Gil et al. 2018), they also provide opportunities for the transmission of parasites and infectious pathogens among associating individuals (Weinstein et al. 2018; Romano et al. 2020; Albery et al. 2021; Cantor et al. 2021; Romano et al. 2021). Wildlife pathogen outbreaks affect most animal taxa, including mammals (Blehert et al. 2009; Fereidouni et al. 2019; Chandler et al. 2021; Kuchipudi et al. 2022), birds (Wille and Barr 2022), amphibians (Scheele et al. 2019), and social insects (Goulson et al. 2015). Weighing the potential risk of infection from social interactions against the benefits of social movements — where to move in relation to other individuals’ positions — is thus a common behavioural context shared by many animal species. Movement strategies incorporating social information — the presence and status of neighbours — can facilitate or reduce spatial associations, and help animals balance the costs and benefits of sociality (Gil et al. 2018; Webber and Vander Wal 2018; Albery et al. 2021; Webber et al. 2022). Animals’ social movements link landscape spatial structure, individual distributions, and the emergent structure of animal societies (Kurvers et al. 2014; Gil et al. 2018; Webber et al. 2022). Together, they influence the dynamics of disease outbreaks in animal populations (Keeling et al. 2001; White et al. 2018*a*; Romano et al. 2020; 2021), and such outbreaks may in turn have cascading effects on landscape structure and community ecology (Monk et al. 2022).

On ecological timescales, pathogen outbreaks often reduce social interactions among individuals. This is due to a combination of mortality-induced decreases in population density (e.g. Fereidouni et al. 2019; Monk et al. 2022), and adaptive behavioural responses by which animals reduce encounters between infected and healthy individuals (Stroeymeyt et al. 2018; Weinstein et al. 2018; Pusceddu et al. 2021; Stockmaier et al. 2021). The latter case includes self-isolating when infected, or avoiding potentially infectious individuals (Stroeymeyt et al. 2018; Weinstein et al. 2018; Pusceddu et al. 2021; Stockmaier et al. 2021). However, when pathogens are first introduced into a population, such as during novel cross-species spillover (Chandler et al. 2021; Kuchipudi et al. 2022), fine-tuned avoidance responses are less likely, as individuals may have no prior experience of cues that indicate infection (Weinstein et al. 2018; Stockmaier et al. 2021). Spreading through host-host contacts, pathogens causing chronic infections (Bastos et al. 2000; Vosloo et al. 2009; Jolles et al. 2021) may instead impose fitness costs, thus selecting against host social behaviour, and hence against social connectivity itself (Altizer et al. 2003; Cantor et al. 2021; Poulin and Filion 2021; Romano et al. 2021; Ashby and Farine 2022).

Yet novel pathogen introductions are primarily studied for their immediate demographic (Fey et al. 2015), and potential medical (Levi et al. 2012; Chandler et al. 2021; Kuchipudi et al. 2022; Wille and Barr 2022) and economic implications (Keeling et al. 2001; Goulson et al. 2015; Jolles et al. 2021), with host evolutionary dynamics (and especially changes in sociality) mostly ignored. This is presumably because the evolution of pathogen host traits, and moreover complex behavioural traits such as sociality, is expected to be slow and not immediately relevant. Since important aspects of animal ecology, including the transmission of foraging tactics (Klump et al. 2021) and migration routes (Guttal and Couzin 2010; Jesmer et al. 2018), depend on social interactions, it is necessary to understand the long-term consequences of pathogen introductions for animal societies. Climate change is only expected to make novel pathogen introductions more common (Sanderson and Alexander 2020; Carlson et al. 2022), making such studies more urgent.

Theory suggests that animal sociality evolves to balance the value of social associations against the risk of pathogen transmission (Bonds et al. 2005; Prado et al. 2009; Ashby and Farine 2022). However, analytical models often reduce animal sociality to single parameters, while it actually emerges from individual decisions conditioned on multiple internal and external cues. Social decision-making and movement often also vary among individuals (Tanner and Jackson 2012; Wolf and Weissing 2012; Spiegel et al. 2017; Gartland et al. 2021), but analytical models are unable to include individual differences in sociability. Epidemiological models based on contact networks can incorporate individual variation in social behaviour by linking these differences to positions in a social network (White et al. 2017; Albery et al. 2020; 2021). Yet network models often cannot capture fine-scale feedbacks between individuals’ social and spatial positions (Albery et al. 2020; 2021), nor spatial variation in infection risk (Albery et al. 2022), making such models sensitive to both the network formation process, and to sampling biases in empirical data collection (White et al. 2017).

Mechanistic, individual-based simulation models (IBMs) suggest themselves as a natural solution; they can incorporate substantial ecological detail, including explicit spatial settings (DeAngelis and Diaz 2019), and detailed disease transmission (White et al. 2018*a*,*b*; Scherer et al. 2020; Lunn et al. 2021). Individual-based models hitherto haved focused on immediate epidemiological outcomes, such as infection persistence, and do not have an evolutionary component (White et al. 2018*b*; Scherer et al. 2020; Lunn et al. 2021). Incorporating an evolutionary component to movement-disease IBMs could allow predictions on important feedbacks between the ecological outcomes of infectious disease and the consequences for the evolution of host behaviour (Cantor et al. 2021). This could include the emergence of tradeoffs in the costs and benefits of sociability (Gartland et al. 2021), with cascading ecological and social effects (Tanner and Jackson 2012; Spiegel et al. 2017; Monk et al. 2022; Webber et al. 2022). The range of animal taxa at risk from a wide array of pathogens and parasites (Sanderson and Alexander 2020; Carlson et al. 2022) makes it important to conceive of models that can capture the key features of diverse host-pathogen dynamics and offer broad conceptual insights (White et al. 2018*a*,*b*).

We built a model that seeks to capture the essential elements of pathogen (or parasite) transmission among animals foraging on patchily distributed resources — this is a common behavioural context shared by many potential host species (White et al. 2018*a*,*b*). We examined the eco-evolutionary consequences of the introduction of a pathogen into a novel host population (such as during cross-species spillover: Bastos et al. 2000; Blehert et al. 2009; Fereidouni et al. 2019; Scheele et al. 2019; Sanderson and Alexander 2020; Carlson et al. 2022; Kuchipudi et al. 2022; Monk et al. 2022; Wille and Barr 2022). In our evolutionary, spatial, individual-based simulation, we modelled the repeated introduction of an infectious pathogen to populations that had already evolved foraging movement strategies in its absence. Our model could be conceived as an abstract representation of, among others, spillovers of foot-and-mouth disease from buffalo to impala (Bastos et al. 2000; Vosloo et al. 2009), or sarcoptic mange from llamas to vicuñas (Monk et al. 2022), current and historic spread of avian influenza among sea-and wading bird species (H5N8 and Related Influenza Viruses 2016; Wille and Barr 2022), or SARS-CoV-2 from humans to deer (Chandler et al. 2021; Kuchipudi et al. 2022).

We compared how social information was used in movement strategies evolved before and after pathogen introduction, and the ecological outcomes for individual intake, movement, and associations with other foragers. Using both IBMs and network epidemiological models (Bailey 1975; White et al. 2017; Stroeymeyt et al. 2018; Wilber et al. 2022), we examined whether pathogen-risk adapted populations were more resilient to the spread of infectious disease than their pathogen-risk naive ancestors. We also investigated the effect of landscape productivity and the cost of infection, which are both expected to influence the selection imposed by pathogen transmission (Hutchings et al. 2000; Almberg et al. 2015; Ezenwa et al. 2016). Overall, we provide a theoretical framework broadly applicable to novel host-pathogen introduction scenarios, and demonstrate the importance of including evolutionary dynamics in movement-disease models.

## Methods

We implemented an individual-based simulation model to represent foraging animals (‘foragers’) seeking discrete, immobile, depleteable food items (see *SI Appendix Fig. S1 – S2*) (Spiegel et al. 2017; Gupte et al. 2021). Food items are distributed over a two-dimensional, continuous-space resource landscape with wrapped boundaries (a torus). Our model, similar to previous eco-evolutionary individual based models (Getz et al. 2015; Gupte et al. 2021; Netz et al. 2021), has two distinct timescales: (1) an ecological timescale comprising of *T* timesteps that make up one generation (*T* = 100 by default), and (2) an evolutionary timescale consisting of 5,000 generations (G). At the ecological timescale, individuals sense local counts of food items and competitors, move according to inherited movement strategies, and forage for food. At the same timescale, individuals that carry an infectious, fitness-reducing pathogen, may, when in close proximity with uninfected individuals, pass on the pathogen with a small probability (see *Pathogen Transmission and Disease Cost*). At the evolutionary timescale, individuals reproduce and transmit their movement strategies (see *Starting Location and Inheritance of Movement Rules*) to the their offspring. The number of offspring is linked both to individuals’ success in finding and consuming food items, and to the duration that they were infected by the pathogen at the ecological timescale. The model was implemented in R and C++ using Rcpp (Eddelbuettel 2013; R Core Team 2020) and the *Boost.Geometry* library for spatial computations (*www.boost.org*); model code is at *github.com/pratikunterwegs/pathomove*.

### Distribution of Food Items

Our landscape of 60 × 60 units contains 1,800 discrete food items, which are clustered around 60 resource ‘kernels’, for a resource density of 0.5 items per unit^2^ (see *SI Appendix Fig. S1 – S2*). This prevents synchronicity in the availability and regeneration of food items. Each available food item can be sensed and harvested by foraging individuals (see below). Once harvested, another food item is regenerated at the same location after a fixed regeneration time R, which is set at 50 timesteps by default; alternative values of 20 and 100 timesteps represent high and low productivity landscapes respectively. Food item regeneration is delinked from population generations. Thus the actual number of available food items is almost always in flux. In our figures and hereafter, we chose to represent R as the number of times a food item would regenerate within the timesteps in a single generation *T* (default = 100), resulting in R values of 1, 2, and 5 for regeneration times of 100, 50 (the default), and 20 timesteps. Items that are not harvested remain on the landscape until they are picked up by a forager. Each food item must be processed, or ‘handled’, by a forager for *T_H_* timesteps (the handling time, default = 5 timesteps) before it can be consumed (Ruxton et al. 1992; Gupte et al. 2021). The handling time dynamic is well known from natural systems in which there is a lag between finding and consuming a food item (Ruxton et al. 1992).

### Individual Foraging and Movement

#### Foraging

Individuals forage in a randomised order, harvesting the first available food item within their movement and sensory range (*d_S_* = *d_M_*, a circle with a radius of 1 unit (see *SI Appendix Fig. S1 – S2*). Once harvested, the item is no longer available to other individuals, leading to exploitation competition among nearby foragers. Furthermore, the location of the item also yields no more cues to other foragers that an item will reappear there, reducing direct cues by which foragers can navigate to profitable clusters of food items. Individuals that harvest a food item must handle it for *T_H_* timesteps (default = 5 timesteps), while all individuals not handling a food item are considered idle (Ruxton et al. 1992; Gupte et al. 2021). As handlers are immobilised at the location where they encountered food, they may be good indirect indicators of the location of a resource cluster (‘social information’) (Danchin et al. 2004; Romano et al. 2020; Gupte et al. 2021). Once individuals finish handling a food item, they return to the non-handling, searching state.

#### Movement

Our model individuals move in small, discrete steps of fixed size (*d_M_* = 1 unit). Each step is chosen based on the individuals’ assessment of local environmental cues, and this assessment is made using evolved movement strategies (as in Gupte et al. 2021; Netz et al. 2021). First, individuals scan their current location, and five equally spaced points around their position, at a distance of 1 unit for three cues (*d_S_*, see *SI Appendix Fig. S1 – S2*): the number of food items (*F*), the number of foragers handling a food item (‘handlers’: *H*) and the number of idle foragers not handling a food item (‘non-handlers’: *N*). Individuals assign a suitability (see Gupte et al. 2021; Netz et al. 2021) to their current position and each of the five locations, using their inherited preferences for each of the cues: *S* = *s_F_F* + *s_H_H* + *s_N_N* + *ε*. The preferences *s_F_*, *s_F_*, and *s_N_* for each of the three cues are heritable from parents to offspring, while *ε* is a very small error term drawn for each location, to break ties among locations. The values of each of the cue preferences *relative to each other* determine individuals’ movement strategies (Gupte et al. 2021). All individuals move simultaneously to the location to which they have assigned the highest suitability (‘step selection’) (akin to step-selection; Fortin et al. 2005); this may be their current location, in which case individuals are stationary for that timestep. Since individuals may differ in their inherited preferences for each of the three cues, two individuals at the same location may make quite different movement decisions based on the same local cues. Handlers, however, are considered immobile and do not make any movement decisions.

### Pathogen Transmission and Disease Cost

We modelled circumstances that are expected to become increasingly common due to rapid global changes; the population evolves for 3/5^th^ of the simulation (until G = 3,000; of 5,000) in the absence of a pathogen, after which a pathogen is introduced in each generation until the end of the simulation (G = 5,000). Our model captures some essential features of pathogen or parasite transmission among animals (White et al. 2017): the pathogen may transmit from infected host individuals to their susceptible neighbours with a per-timestep probability *p* of 0.05. This transmission is only possible when the two individuals are within a the transmission distance, *d_β_*. For simplicity, we set *d_β_* to be the movement range (1 unit). Once transmitted, the pathogen is assumed to cause a chronic disease which reduces host energy stores by a fixed amount called *δE* in every following timestep; *δE* is set to 0.25 by default (alternative values: 0.1, 0.5). Since novel pathogen introductions can periodically re-occur in natural environments (Bastos et al. 2000; Vosloo et al. 2009; Almberg et al. 2015; Goulson et al. 2015; Jolles et al. 2021; Carlson et al. 2022; Wille and Barr 2022), we set up our model such that the pathogen was introduced to 4% of individuals in each generation (N = 20; ‘primary infections’). This is necessary to kick-start the pathogen-movement eco-evolutionary feedback dynamics, and populations may indeed repeatedly acquire novel pathogens (or strains) through external sources, such as infected individuals of other spatially overlapping species (e.g. Bastos et al. 2000; Keeling et al. 2001; Vosloo et al. 2009; Chandler et al. 2021; Carlson et al. 2022; Kuchipudi et al. 2022; Monk et al. 2022; Wille and Barr 2022). For completeness, we also considered scenarios in which novel pathogen introductions only occur sporadically in the generations after the initial event, rather than in every generation (see *SI Appendix*).

### Starting Location and Inheritance of Movement Rules

For simplicity, we considered a population of haploid individuals with discrete, non-overlapping generations, and asexual inheritance. At the end of the parental generation, the net lifetime energy of each individual was determined as the difference of the total energy gained through food intake and the energy lost through infection. In the *SI Appendix,* we also consider an alternative implementation in which potential immune resistance against the pathogen requires a certain percentage of individual intake, reducing the value of each food item. The parental population produces an offspring population (of the same size) as follows: to each offspring, a parent is assigned at random by a weighted lottery, with weights proportional to lifetime net energy (an algorithm following the replicator equation) (Hofbauer and Sigmund 1988; Hamblin 2013). This way, the expected number of offspring produced by a parent is proportional to the parent’s lifetime success (Hofbauer and Sigmund 1988). The movement decision-making cue preferences *s_F_*, *s_H_*, and *s_N_* are subject to independent random mutations with a probability of 0.01. The mutational step size (either positive or negative) is drawn from a Cauchy distribution with a scale of 0.01 centred on zero. Thus, while the majority of mutations are small, there can be a small number of very large mutations. As in real ecological systems, individuals in the new generation are intialised around the location of their parent (within a standard deviation of 2.0), and thus successful parents give rise to local clusters of offspring (see an alternative implementation in *SI Appendix*).

### Model Output

#### Social Information Use

To understand the evolution of movement strategies, and especially how individuals weighed social information, we recorded the population’s evolved cue preferences in every second generation, and interpreted them using the ‘behavioural hypervolume’ approach (Bastille-Rousseau and Wittemyer 2019). We classified individuals based on how they used social information — the presence and status of competing foragers — into four social movement classes: (1) agent avoiding, if *s_H_*, *s_N_* < 0, (2) agent tracking, if both *s_H_*, *s_N_* > 0, (3) handler tracking, if *s_H_* > 0, *s_N_* < 0, and (4) non-handler tracking, if *s_H_* < 0, *s_N_* > 0. We calculated the relative importance of social cues — *H*, *N* — to each individual’s movement strategy as *SI_imp_* = (|*s_H_*| + |*s_N_*|)/(|*s_H_*| + |*s_N_*| + |*s_F_*|), with higher values indicating a greater importance of social cues.

#### Proximity-Based Social Network

We created a proximity-based adjacency matrix by counting the number of times each individual was within the sensory and pathogen transmission distance *d_β_* (= *d_S_*, *d_M_* = 1 unit) of another individual (Whitehead 2008; Wilber et al. 2022). We transformed this matrix into an undirected social network weighted by the number of pairwise encounters: in a pairwise encounter, both individuals were considered to have associated with each other (White et al. 2017). The strength of the connection between any pair was the number of times the pair were within *d_β_* of each other over their lifetime. We logged encounters and constructed social networks after every 10% of the total generations (i.e., every 500^th^ generation), and at the end of the simulation, and omitted ephemeral pairwise associations with a weight *<* 5.

### Model Analysis

We plotted the mix of social information-based movement strategies evolved across generations in each parameter combination. Focusing on our default scenario (*δE* = 0.25, R = 2), we visualised the mean per-capita distance moved, mean per-capita intake, and mean per-capita encounters with other foragers. We examined how the three main social movement strategies — agent avoidance, agent tracking, and handler tracking — changed in frequency over generations. We also examined differences among strategies in the movement distance, associations with other agents, and frequency of infection, after they had reached an eco-evolutionary equilibrium following pathogen introduction (G > 3,500). We visualised the proximity based social networks of populations in a representative scenario (*δE* = 0.25, R = 2), focusing on the generations just before and after the pathogen introduction events begin (pre-introduction: G = 3,000; post-introduction: G = 3,500). We plotted the numbers of individuals infected in each generation after pathogen introduction to examine whether evolutionary changes in movement strategies actually reduced infection spread. We also ran simple network epidemiological models on the emergent individual networks in generations 3,000 and 3,500 (Bailey 1975; White et al. 2017; Stroeymeyt et al. 2018; Wilber et al. 2022), for robust comparisons of potential pathogen spread in pathogen-risk naive and pathogen-risk adapted populations, respectively.

### Data and Code Availability

The *Pathomove* simulation model code is available on Zenodo at https://zenodo.org/record/6782640, and on Github at github.com/pratikunterwegs/pathomove. Code to run the simulations and analyse the output is on Zenodo at https://zenodo.org/record/6782665, and on Github at: github.com/pratikunterwegs/patho-move-evol.

## Results

In our model, individuals move and forage on a landscape with patchily distributed food items, and select where next to move in their vicinity, based on inherited preferences for environmental cues — food items, and other individuals (see *SI Appendix Fig. S1*). Food items, once consumed, regenerate at a rate R, and pathogen infection imposes a per-timestep cost *δE*. We classified individuals’ social movement strategies in our model using a simplified ‘behavioural hypervolume’ approach (Bastille-Rousseau and Wittemyer 2019), based on the sign of their preferences for successful foragers handling a food item (‘handlers’, preference *s_H_*), and for unsuccessful foragers still searching for food (‘non-handlers’, preference *s_N_*). In our default scenario, R = 2, food regenerates twice per generation, and *δE* = 0.25, i.e., consuming 1 food item offsets 4 timesteps of infection. Over the 3,000 generations before the introduction of the pathogen, populations reached an eco-evolutionary equilibrium where the commonest social movement strategy was to prefer moving towards both handlers and non-handlers (‘agent tracking’; *s_H_*, *s_N_* > 0; but see below) (Fig. 1A).

**Figure 1:**
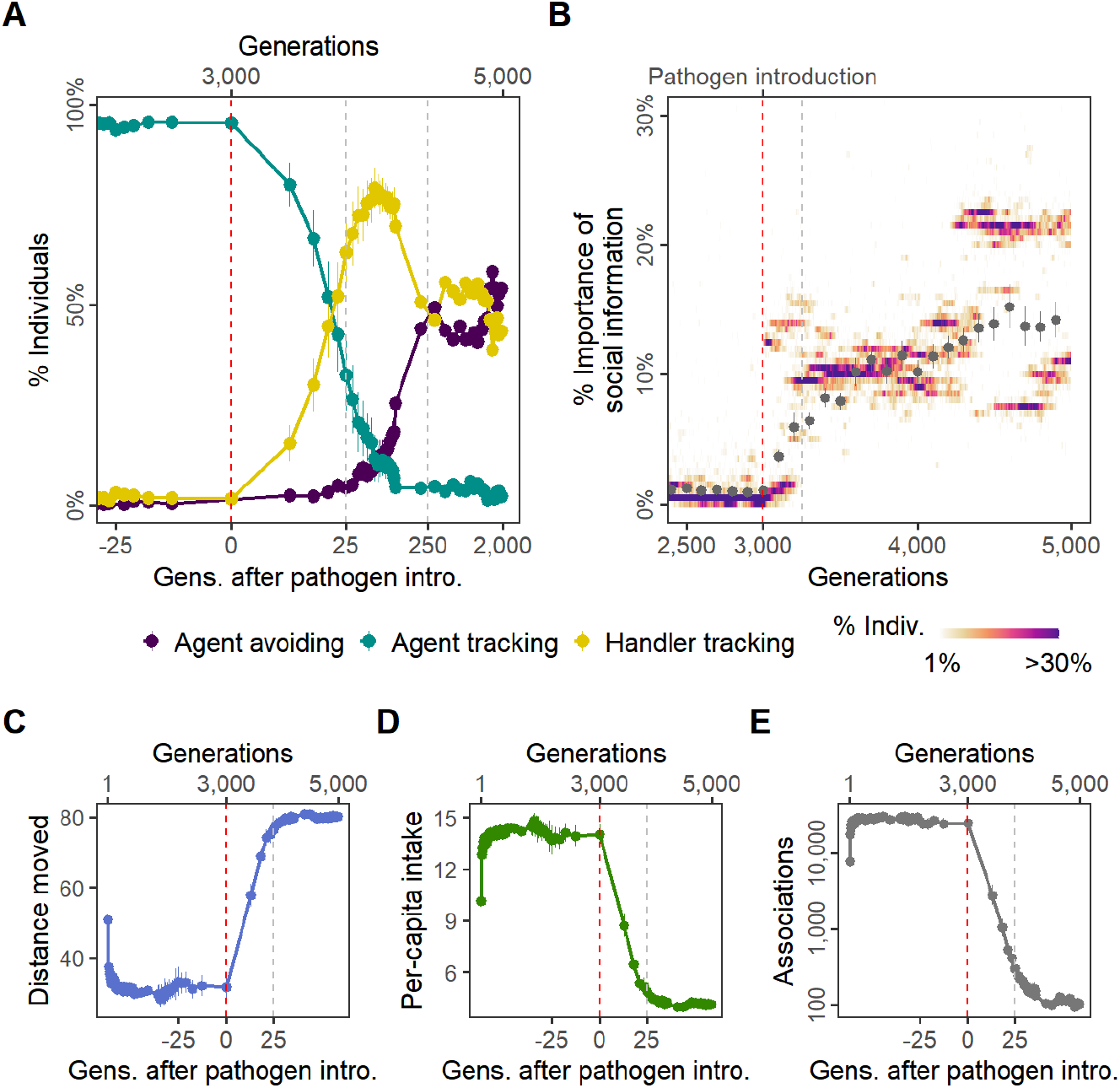
Pathogen introduction leads to rapid evolutionary changes in social information use, with cascading effects on population ecological outcomes. **(A)** Before pathogen introduction in the default scenario (R = 2, *δE* = 0.25), populations rapidly evolve a social movement strategy that tracks all other individuals (‘agent tracking’; *G* ≤ 3,000) — however, their overall movement strategy is primarily guided by the presence of food items **(B)**. Pathogen introduction leads to the rapid replacement, within 25 generations, of agent tracking with ‘handler tracking’ (preference for successful foragers; 3,000 *G* 3,025). Within 250 generations, ‘agent avoidance’ (avoidance of both successful and unsuccessful foragers; *G* > 3,250) also becomes common, stably co-existing with the handler tracking strategy in an eco-evolutionary equilibrium. **(B)** After pathogen introduction (*G* > 3,000), the importance of social cues (the presence of other individuals; the sum of the absolute, normalised preferences *sH, sN*) increases substantially on average (grey points). Additionally, there is significant variation in the importance of social cues to individuals (shaded regions), which is not captured by the mean or standard error. At G = 4,500, for example, social information comprises ≈ 10% of some individuals’ movement strategies, but some individuals have evolved a stronger weight for social cues (> 20%). The rapid change in social movement strategies following pathogen introduction has cascading effects on ecological outcomes. Individuals, which have evolved strong aversions to at least some kinds of foragers (depending on their strategy), **(C)** move more on average, **(D)** have only 25% of the pre-pathogen average intake, and **(E)** have 100-fold fewer associations with other individuals. All panels show data averaged over 10 replicates, but shaded region in panel B shows only a single replicate for clarity.

### Rapid Evolutionary Shift in Social Movement Strategies Following Pathogen Introduction

Introducing an infectious pathogen to 4% (n = 20) of individuals in each generation (after G = 3,000), leads to a remarkably rapid evolutionary shift — within only 25 generations of pathogen introduction — in how social information is incorporated into individuals’ movement strategies. There is a marked increase in the frequency of individuals that track successful foragers, but avoid non-handlers (‘handler tracking’; *s_H_* > 0, but *s_N_* < 0) (Fig. 1A; 3,000 < *G* < 3,025). Surprisingly, after a brief period (in evolutionary terms) of handler tracking being the most common strategy, a third strategy also becomes more common: avoiding both handlers and non-handlers (‘agent avoiding’; *s_H_*, *s_N_* < 0). Within 250 generations after pathogen introduction, agent avoiding becomes as common as the handler tracking strategy, and this appears to be a stable equilibrium that is maintained until the end of the simulation (2,000 generations after pathogen introduction; Fig. 1A). The *SI Appendix* shows how the occurrence of rapid evolutionary shifts is broadly robust to modelling assumptions; in brief, such shifts occur even when individuals cannot benefit from evolved adaptation to local conditions (Badyaev and Uller 2009), and when the pathogen saps a percentage, rather than an absolute value, from daily intake.

In addition to qualitative changes in social movement strategies, pathogen introduction also leads to social information becoming more important to movement decisions. Prior to pathogen introduction (*G* < 3,000), individuals’ handler-and non-handler preferences (|*s_H_*| + |*s_N_*|; taken together, social information) barely influence their movement strategies (Fig. 1B). These are instead guided primarily by the preference for food items (*s_F_*; see *Model and Analysis*; see also *Supplementary Material Fig. 1*). Social movement decisions are joint outcomes of individual preferences for social cues and the cue value: consequently, in clustered populations (see below), even small positive values of *s_H_* and *s_N_* lead to strong emergent sociality. After pathogen introduction, there is a substantial increase in the average importance of individuals’ preferences (or aversions) for the presence of other foragers (Fig. 1B). However, there is significant variation among individuals in the importance of social information to their movement strategies, with distinct evolved polymorphisms that vary substantially between simulation replicates (Fig. 1B).

### Disease-dominated Ecological Cascade Due to Evolutionary Shift in Movement Strategies

The evolutionary shift in social movement strategies causes a drastic change in ecological outcomes (Fig. 1C – E; see *SI Appendix Fig. S3* for other scenarios). There is a sharp increase in mean distance moved by individuals; while pre-introduction individuals moved 35% of their lifetimes on average (i.e., 35 timesteps; handling for the remainder), post-introduction, individuals move for 80% of their lifetimes (i.e., 80 timesteps; Fig. 1C). The handler tracking and agent avoiding strategies lead individuals to move away from groups of individuals (‘dynamic social distancing’; Pusceddu et al. 2021). Individuals being most likely to be found near resource clusters, this leads to movement away from productive areas of the landscape. Consequently, there is a rapid, four-fold drop in mean per-capita intake after pathogen introduction (Fig. 1D). The concurrent, near 100-fold drop in encounters between individuals after pathogen introduction (Fig. 1E) suggests that most encounters were likely taking place on or near resource clusters. The reductions in intake observed are equivalent to those expected from halving landscape productivity (*SI Appendix Fig. S3*). Our model shows how even a non-fatal pathogen, by influencing the evolution of movement strategies, can have substantial indirect ecological effects — a disease dominated ecological cascade (Monk et al. 2022).

### Co-existence of Social Movement Strategies

At eco-evolutionary equilibrium (G > 3,500) the relationship between movement and avoiding associations (and further, infection) is mediated by individual differences in how exactly social information is incorporated into movement strategies. Individuals using the agent avoiding strategy move more than handler tracking ones (Fig. 2A), about 85% of their lifetime (default scenario: R = 2; *δE* = 0.25). At this limit, every step moved allows them to avoid approximately 2 encounters with other individuals. Handler tracking individuals move much less (~ 60% – 80%), but are able to avoid approximately 20 encounters with other individuals with every extra step. These differences may explain why agent avoiding and handler tracking individuals have similar mean infection rates, at ~25% and ~33% respectively (Fig. 2B). All other strategies, especially the agent tracking strategy common in pre-introduction populations, are barely able to translate increased movement into fewer associations (Fig. 2A). These strategies have a wide range of infection rates (Fig. 2B), potentially because they are very rare — these likely represent mutants that do not give rise to persistent lineages.

**Figure 2:**
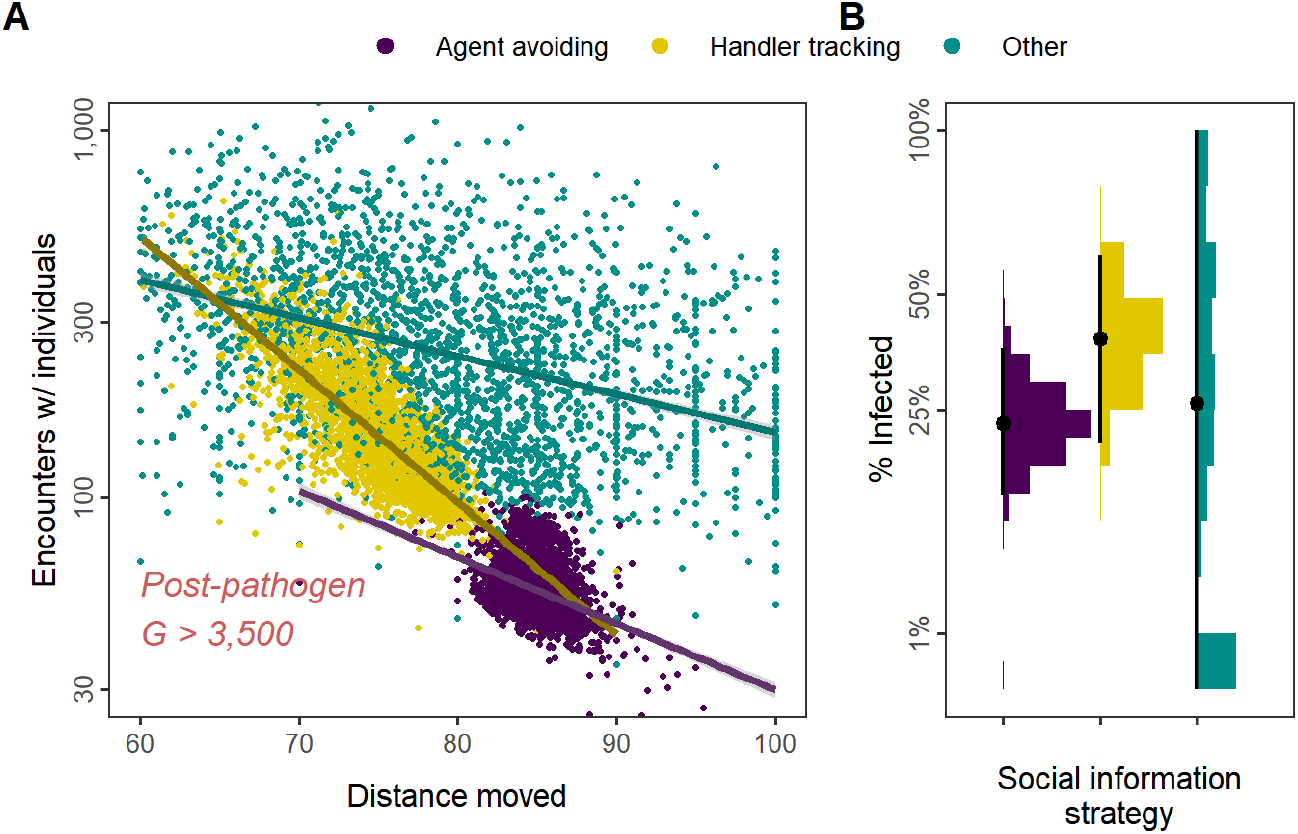
Social movement strategies trade movement for associations through dynamic social distancing, leading to differences in infection rates. In post-introduction populations at eco-evolutionary equilibrium (G > 3,500), **(A)** both agent avoiding and handler tracking individuals can reduce encounters with other individuals by moving to avoid other foragers (dynamic social distancing). Handler tracking individuals have many more encounters than agent avoiding individuals, but surprisingly, are better able to reduce encounters through increased movement. Individuals using other strategies (mostly agent tracking) have a wider range of movement distances, but cannot efficiently avoid other foragers by moving more. **(B)** Avoiding all other foragers leads to marginally lower infection rates than tracking successful foragers (and avoiding unsuccessful ones; handler tracking). Surprisingly, rare pre-introduction strategies such as following any nearby individuals (agent tracking) may also have low infection rates, potentially due to their rarity. Panel A shows linear model fits with a log scale Y-axis; panel B shows infection rates; all data represent generation-and replicate-specific means (G > 3,500; R = 2, *δE* = 0.25).

### Reorganisation of Spatial-social Structure

Following pathogen introduction, the mixture of individual-level movement strategies elicits a substantial re-organisation of emergent spatial and social structure at the population level. Pre-introduction populations are strongly clustered in space (Fig. 3A), due to movement strategies that favour following most other foragers. This spatial proximity means that most individuals encounter each other at least once, leading to numerous unique partners (the ‘degree’) for each forager (Fig. 3 inset 1: *blue*). In con-trast, the spread-out networks in pathogen-risk adapted populations suggest that most foragers move substantially from their initial locations over their lifetime, associating only ephemerally with foragers from all over the landscape (Fig. 3B). This reflects movement strategies which lead to near-perpetual movement to avoid associations; a sort of dynamic social distancing seen in real animal societies under risk of pathogen spread (Stroeymeyt et al. 2018; Weinstein et al. 2018; Pusceddu et al. 2021; Stockmaier et al. 2021). This dispersed population structure means that most pathogen-risk adapted foragers encounter fewer than 10% of the population over their lifetime (Fig. 3 inset 1: *red*).

**Figure 3:**
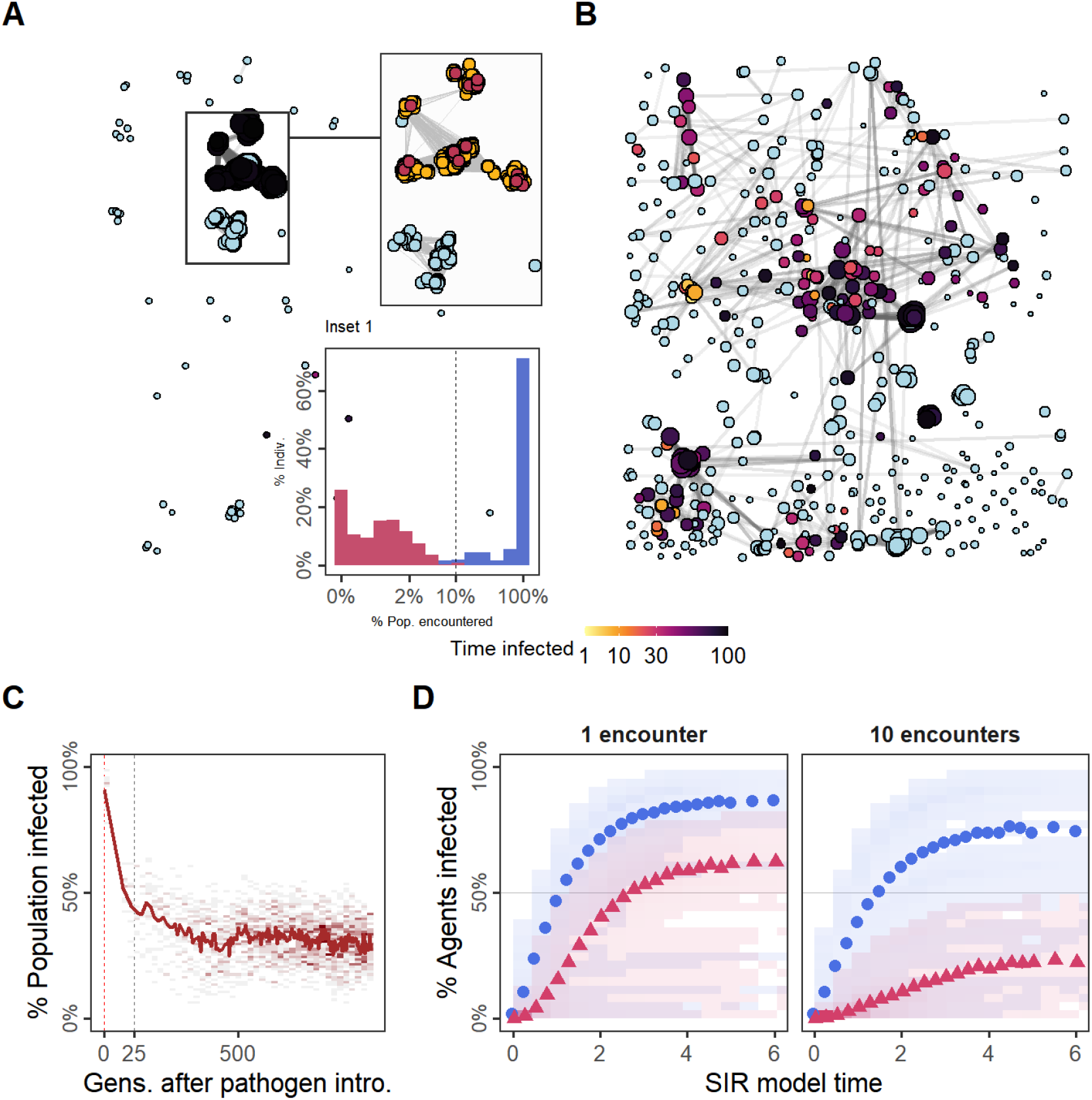
Reduced spatial-social clustering and disease transmission in populations adapted to the presence of an infectious pathogen. pathogen-risk naive populations (**A**; G = 3,000) are much more spatially clustered than pathogen-risk adapted populations (**B**; G = 3,500), and are thus rapidly infected (red: primary infections; yellow: secondary infections; blue: never infected). Pre-introduction individuals encounter many more unique neighbours (**inset 1**, blue) than pathogen-risk adapted individuals (**inset 1**; red). Dashed grey line represents 10% of individuals encountered (N = 50). Main panels show social networks from a single replicate of the default scenario (R = 2, *δE* = 0.25), insets show 10 replicates. Nodes represent individuals positioned at their final location. Connections represent pairwise encounters, and node size represents encounters (larger = more encounters). Darker node colours indicate longer infection (light blue = no infection). **(C)** In the first generations following pathogen introduction, nearly every single individual in the population is infected. However, within 25 generations, tracking the evolutionary shift towards movement strategies that avoid some or all other individuals, only about 50% of individuals are ever infected; this drops to a stable 30% within 500 generations after pathogen introduction. **(D)** The progression of two hypothetical diseases, requiring a single encounter, or 10 encounters for a potential transmission, on emergent social networks. The transmission of both diseases is reduced in populations with disease-adapted movement strategies (pre-introduction: G = 3,000, blue circles; post-introduction: G = 3,500, red triangles). Subfigures in panel D show means of 25 SIR model replicates (transmission rate *β* = 5.0, recovery rate *γ* = 1.0), run on emergent social network; both panels represent 10 simulation replicates the default scenario.

### Pathogen-risk adapted Movement Strategies Make Animal Societies More Resilient to the Spread of Disease

Nearly every individual in the generations just after pathogen introduction was infected. However, tracking the evolutionary change in movement strategies, the number of infected individuals fell to just about 50% within 25 generations (Fig. 3C). To examine potential pathogen spread in pre-introduction populations, we ran a simple epidemiological model on the social networks emerging from individuals’ movements before and after pathogen introduction (pre-introduction: G = 3,000; post-introduction: G = 3,500). We modelled two diseases, (*i*) first, a disease requiring one encounter, and (*ii*) second, a disease requiring ten encounters between individuals for a potential transmission event (transmission rate *β* = 5.0, recovery rate *γ* = 1.0).

Both the single encounter and multiple encounter diseases would infect 75% – 80% of individuals when spreading through the networks of pre-introduction populations (Fig. 3D) pathogen-risk adapted populations’ social networks are more resilient to both the single encounter and multiple encounter disease, compared to their pre-introduction, pathogen-risk naive ancestors (Fig. 3D), as these social networks are sparser and individuals are more weakly connected (Fig. 3D; see Fig. 3D). Less than 60% of post-introduction populations were finally infected by the single encounter disease, compared with *>* 75% of pre-introduction, pathogen-risk naive ancestors; in pathogen-risk adapted populations, the spread of the multiple encounter disease was even slower (ever infected: ≈ 20%).

### Usefulness of Social Information and Infection Cost Influence Evolution of Social Movement Strategies

We further explored the effect of two ecological parameters, landscape productivity (*R* ∈ 1, 2, 5) and infection cost per timestep (*δE* ∈ 0.1,0.25,0.5) on simulation outcomes. Before pathogen introduction, landscape productivity alone determines the value of social information, and thus which social movement strategies evolve (Fig. 4). On low-productivity landscapes (R = 1), social information is valuable as direct resource cues are scarce; here, the handler-tracking strategy persists. On high-productivity landscapes (*R* ∈ 2, 5), social information is less valuable as individuals can directly detect food items more often; here, the agent tracking strategy is most common. Across parameter combinations, the introduction of the infectious pathogen leads to a rapid evolutionary shift in social movement strategies. The benefits of social information, and infection cost jointly determine how pathogen introduction alters the mix of social movement strategies, but populations generally shift away from indiscriminate agent tracking, as that strategy is associated with higher infection risk (see Fig. 3A).

**Figure 4:**
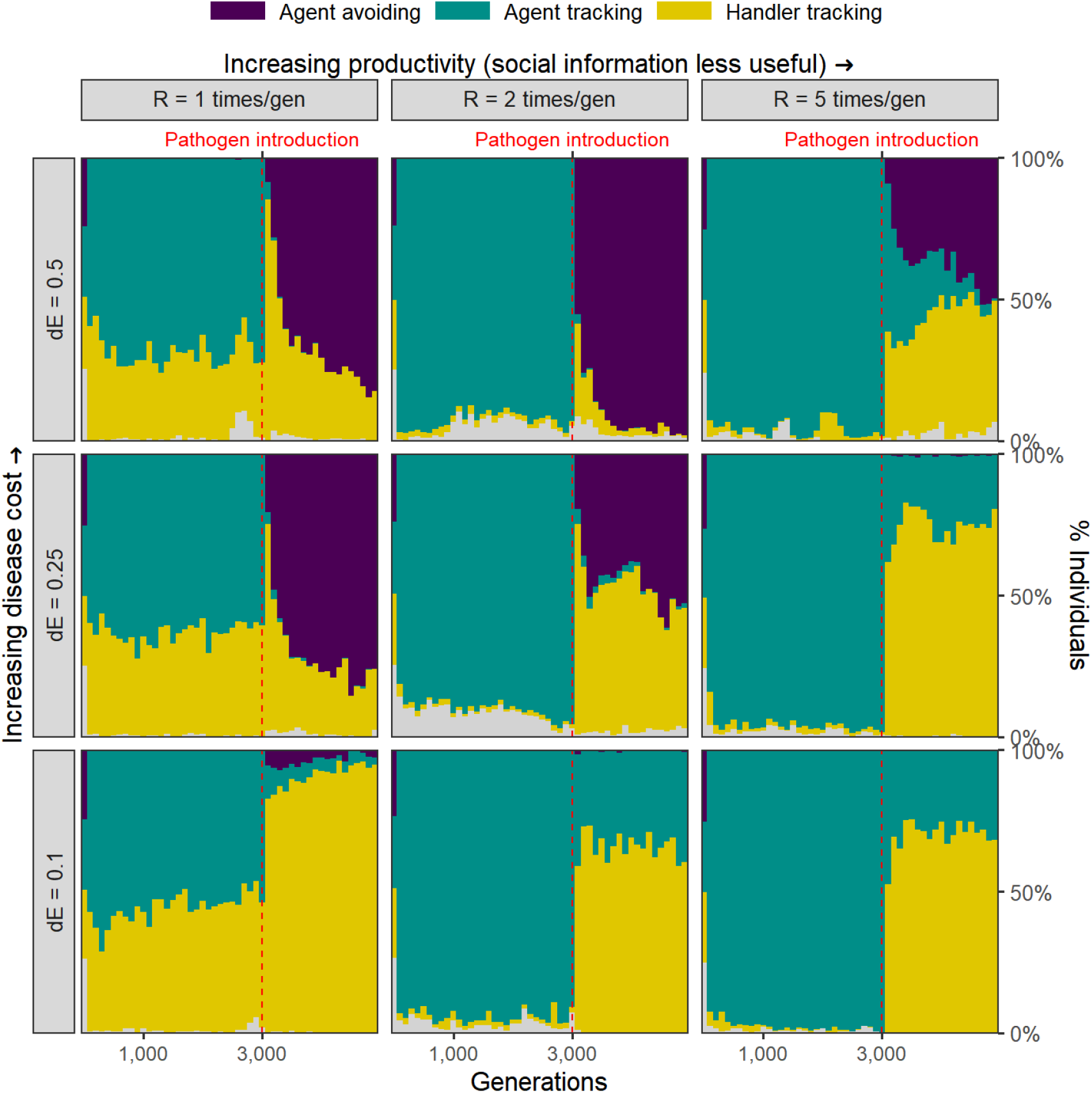
The balance of infection cost and the usefulness of social information together shape the rapid evolutionary change in movement strategies triggered by pathogen introduction. Pre-introduction (G = 3,000; dashed line) populations contain a mix of individuals that either track all foragers (agent tracking), or only successful foragers (handler tracking). Handler tracking is more common on low-productivity landscapes (R = 1), where social information is more useful to find patchily distributed resources. After pathogen introduction, handler tracking rapidly becomes the most common strategy when the apparent usefulness of social information is greater than the cost of infection. This occurs both when productivity is low (R = 1) and infection costs are low (*δE* = 0.1), but also when productivity is high (R = 5) with intermediate infection costs (*δE* = 0.25). When the cost of infection outweighs the apparent usefulness of social information, the agent avoidance (avoiding both successful and unsuccessful foragers) emerges and rapidly becomes a common strategy (*δE* = 0.5; *δE* = 0.25, R = 1). In scenarios of high landscape productivity combined with low infection costs (e.g. R = 5, *δE* = 0.1), the agent tracking strategy persists in a large proportion after pathogen introduction, as these individuals can balance disease costs with intake alone. All panels show mean frequencies over 10 replicate simulations in 100 generation bins; frequencies are stacked. Grey areas show the relatively uncommon ‘non-handler’ tracking strategy.

When the benefit of social information is equivalent to the cost of infection, the handler tracking strategy is common (R =1, *δE* = 0.1; R = 5, *δE* = 0.25). When apparent social information benefits are lower than infection costs (e.g. *δE* = 0.5), the agent avoiding strategy is common. The effect of landscape productivity in obviating a sensitivity to social information cues (especially, conspecific status) is also eroded by pathogen introduction. On high-productivity landscapes where individuals were indiscriminately social, (*R* ∈ 2, 5, *δE* = 0.1), the handler tracking strategy becomes common, as individuals prioritise higher-quality social information (handlers, which indicate a resource cluster). However, high landscape productivity can also compensate for the cost of infection, as evidenced by the agent tracking strategy remaining prevalent: this is only possible if these individuals can consume sufficient resources to overcome disease costs.

## Discussion

Our general model captures important features of infectious pathogen (or parasite) transmission among host animals in a (foraging) context that is relevant to most species. The combination of ecological, evolutionary, and epidemiological dynamics in a spatial setting is unprecedented for movement-disease models, and extends current understanding of animal spatial and social ecology (Kurvers et al. 2014; Webber and Vander Wal 2018; Romano et al. 2020; Albery et al. 2021; Romano et al. 2021; Webber et al. 2022). Presently, most movement-disease models are non-evolutionary (White et al. 2017; 2018*b*; Scherer et al. 2020; Lunn et al. 2021), presumably because evolution is expected to be too slow to impact epidemiological-ecological outcomes (Monk et al. 2022). We demonstrate the pitfalls of this assumption: evolutionary transitions in sociality occur over fewer generations than required for the development of key aspects of animal ecology, such as migration routes (Jesmer et al. 2018; Cantor et al. 2021). We also demonstrate the tension inherent to sociality under the risk of an infectious pathogen, in an explicitly spatial context. Our work shows how qualitatively and quantitatively different social movement strategies — making different trade-offs between social information and infection risk — can co-exist in a single population (Wolf and Weissing 2012; Webber and Vander Wal 2018; Gartland et al. 2021; Webber et al. 2022).

Prior to pathogen introduction, the value of social information influenced which social movement strategies were evolved. Individuals initialised (‘born’) near their parent’s final location may benefit from ‘ecological inheritance’ (Badyaev and Uller 2009) of their parent’s favourable position near resource clusters (see *SI Appendix Fig. S2, S4*). Avoiding potential competitors (and kin) thus correlates with avoiding profitable areas, and this leads to the persistence of the indiscriminately social agent tracking strategy, despite the evident costs of exploitation competition. In an alternative implementation with large-scale natal dispersal, handler tracking is the commonest strategy prior to pathogen introduction (see *SI Appendix*). Following pathogen introduction, the agent tracking strategy of our default scenario allows the disease to spread very easily among entire lineages of social individuals (see Fig. 3A) (Kurvers et al. 2014). This neatly demonstrates why the risk of infection or parasitism could be among the mechanisms underlying density dependence in natal dispersal decisions (Travis et al. 1999).

Following pathogen introduction, the evolutionary shift in social movement strategies is much more rapid than the timescales usually associated with the evolution of complex traits such as sociality (about 25 generations). Avoiding potentially infectious individuals is a key component of navigating the ‘landscape of disgust’ (Weinstein et al. 2018). Our results show that sensitivity to cues of high pathogen transmission risk can rapidly evolve following the introduction of a novel pathogen, with a complete replacement of the hitherto dominant social strategy. The emergence of qualitative individual variation in social movement strategies, and especially the trade-off between movement, associations, and infection risk also demonstrates the evolution of ‘sociability as a personality trait’ (Gartland et al. 2021). We also find substantial individual variation in the quantitative importance of social cues overall, which is a key component of the evolution of large-scale collective behaviours, such as migration (Guttal and Couzin 2010). Our work suggests how, by leading to the necessary diversity in social movement strategies, a novel pathogen may actually lay the groundwork for the evolution of more complex collective behaviour. Nonetheless, the rapid decreases in social interactions should primarily prompt concern that the evolutionary consequences of pathogen introduction could slow the transmission of, and erode, animal culture (Cantor et al. 2021) — including foraging (Klump et al. 2021) and migration behaviours (Guttal and Couzin 2010; Jesmer et al. 2018).

In our model, landscape productivity (R), is a proxy for the usefulness of sociality overall, as social information is less useful when direct resource cues are abundant (high R). Social information benefits in disease models often have no mechanistic relationship with the subject of the information (e.g. food or predators) (Ashby and Farine 2022). In contrast, social information benefits in our model are emergent outcomes of animal movement and foraging behaviour. Our predictions may help explain intra-and inter-specific diversity in social systems across gradients of infection risk and the usefulness of social information (Altizer et al. 2003; Sah et al. 2018), and studies tracking social movements and potential for disease spread could form initial tests of our basic predictions (Wilber et al. 2022). While our individuals do not die, the evolved pathogen-risk adapted, dynamic social distancing strategies (Stockmaier et al. 2021) lead to a significant worsening (equivalent to a halving) of individuals’ intake. In real systems, this could increase populations’ susceptibility to extreme climate change related mortal-ity events (Fey et al. 2015).

More positively, animals may be able to adapt relatively quickly to the spillover and eventual persistence of infectious pathogens, even when they cannot specifically detect and avoid infected individuals (Altizer et al. 2003; Stroeymeyt et al. 2018; Pusceddu et al. 2021; Stockmaier et al. 2021). While the most noticeable effect of pathogen outbreaks is mass mortality (Fey et al. 2015), even quite serious pathogens — Sarcoptic mange (Almberg et al. 2015), foot-and-mouth disease (Bastos et al. 2000; Vosloo et al. 2009; Jolles et al. 2021), SARS-CoV-2 (Chandler et al. 2021; Kuchipudi et al. 2022), and avian influenza (H5N8 and Related Influenza Viruses 2016; Wille and Barr 2022) among others — appear to spread at sub-lethal levels for many years between lethal outbreaks. Our model shows how disease-dominated ecological cascades (Monk et al. 2022) could occur even without mortality effects, due to evolutionary shifts in sociality alone. The altered ecological state (here, less resource consumption, as in Monk et al. 2022) may be maintained long after — and indeed because — a population has adapted to be less social in the presence of a pathogen. Our work suggests that decreased sociality resulting from adaptation to a novel pathogen could slow the transmission of future novel pathogens. While decreased sociality could also reduce the prevalence of previously endemic pathogens adapted to a more social host, it may also degrade ‘social immunity’ through reduced sharing of beneficial commensal microbes, or of low, immunising doses of pathogens (Almberg et al. 2015; Ezenwa et al. 2016).

Our model results are contingent upon sustained introduction of the pathogen (or its novel strains) to host populations. More sporadic introductions (once every few generations) apparently do not cause evolutionary shifts in social movement (*SI Appendix*). Yet repeated pathogen and parasite introductions among susceptible populations appear to be quite common (Bastos et al. 2000; Vosloo et al. 2009; Levi et al. 2012; H5N8 and Related Influenza Viruses 2016; Scherer et al. 2020; Jolles et al. 2021; Wille and Barr 2022). Such introductions are often detected only among easily observed groups such as birds (Wille and Barr 2022), or after evident mass mortality events (Fey et al. 2015; Fereidouni et al. 2019). Seasonal host-pathogen dynamics could and do keep pathogens circulating in reservoir hosts, with regular pulses in primary infections similar to our model (e.g. due to new calves in African buffalo hosting foot-and-mouth disease: Jolles et al. 2021, or winter peaks in mange among wolves: Almberg et al. 2015). Existing host-pathogen dynamics, and potential pathogen range expansions, could thus provide more frequent opportunities for novel transmissions to overlapping species than previously guessed. Our model shows how this provides a powerful selective force in favour of detecting and avoiding infection risk cues (Weinstein et al. 2018).

In order to be widely applicable to diverse novel host-pathogen introduction scenarios, our model is necessarily quite general. A wide diversity of pathogens and their dynamics remains to be accurately represented in individual-based models (White et al. 2017; 2018*b*; Scherer et al. 2020; Lunn et al. 2021). Our framework can be expanded and specifically tailored to real-world situations in which populations are repeatedly exposed to novel pathogens (or strains) (Bastos et al. 2000; Scherer et al. 2020; Chandler et al. 2021; Jolles et al. 2021; Kuchipudi et al. 2022; Wille and Barr 2022). Such detailed implementations could include aspects of the pathogen life-cycle (White et al. 2017; 2018*a*), account for sociality as a counter to infection costs (Almberg et al. 2015; Ezenwa et al. 2016), or model host-pathogen sociality-virulence co-evolution (Bonds et al. 2005; Prado et al. 2009; Ashby and Farine 2022). Future work would ideally combine wildlife monitoring and movement tracking across gradients of pathogen prevalence, to detect novel cross-species spillovers (Chandler et al. 2021; Kuchipudi et al. 2022) and study the spatial and epidemiological consequently of animal movement strategies (Bastille-Rousseau and Wittemyer 2019; Monk et al. 2022; Wilber et al. 2022). Our model shows why it is important to consider evolutionary responses in movement-disease studies, and provides a general framework to further the integration of evolutionary approaches in wildlife spatial epidemiology.

## Acknowledgements

We thank Jan Kreider for helpful feedback on an early draft of the manuscript; and Thijs Janssen for help with the simulation model code. We thank the Center for Information Technology of the University of Groningen for providing access to the *Peregrine* high performance computing cluster to run simulations. P.R.G was supported by an Adaptive Life Programme grant made possible by the Groningen Institute for Evolutionary Life Sciences (GELIFES). J.G. was supported by a grand from the Netherlands Organization for Scientific Research (NWO-ALW; ALWOP.668). F.J.W. and P.R.G acknowledge funding from the European Research Council (ERC Advanced Grant No. 789240).

