## Supplementary material for "Novel pathogen introduction rapidly alters evolved movement strategies, restructuring animal societies": SI Appendix

### Supplementary Material for *Novel pathogen introduction rapidly alters the evolution of movement, restructuring animal societies*

Pratik R. Gupte  Gregory F. Albery  Jakob R. L. Gismann  Amy Sweeny
Franz J. Weissing

2022-07-03

**4 References** **11**

### 1 Model description

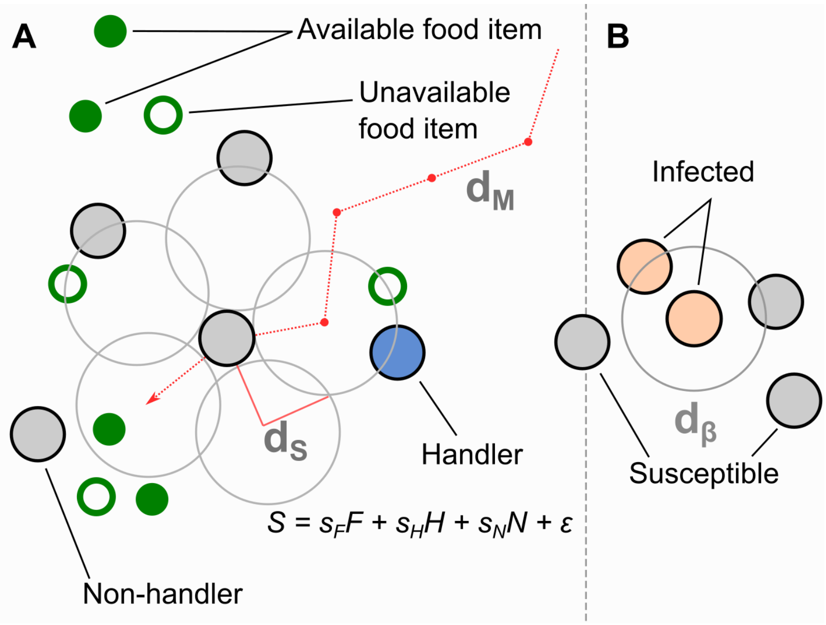

**Figure S1: Model implementation of discrete movement steps in continuous space, with movement steps selected based on inherited preferences for environmental cues.** In our model, **(A)** individuals search for food items (**green circles**), which may be immediately available (**filled green circles**; $F$), or may be available only in the future (**open green circles**). Individuals can sense only available items, and not unavailable ones. However, given our landscape structure, food items are clustered, making available items a good indicator of where resource clusters are (see next figure). Individuals can also sense other foraging individuals, and can sense whether they have successfully found, and are handling, a food item (handlers; **blue circles**), or whether they are unsuccessful foragers still searching for food (non-handlers; **filled grey circles**; $N$). To decide where to move, individuals sample their environment for these three cues (*F, H, N*) at 5 locations around themselves (**large open grey circles**), and have a sensory range of $d_S$. When the sensory range is relatively large (default = 1.0 units), there is some small overlap in samples. Individuals assign each potential direction a *suitability*, $S = s_F F + s_H H + s_N N + \epsilon$, where the coefficients $s_F, s_H, s_N$ are inherited preferences for environmental cues, and $\epsilon$ is a small error term that helps break ties between locations. In our implementation, the sensory distance ($d_S$) and the movement distance ($d_M$) are the same, 1.0 units. **(B)** Our infectious pathogen is transmitted between infected (**orange circles**) and susceptible (**filled grey circles**) individuals, with a probability $p$ = 0.05, when they are within a distance $d_\beta$ of each other. In our implementation, $d_\beta$ is the same as $d_S, d_M$ = 1.0 units.

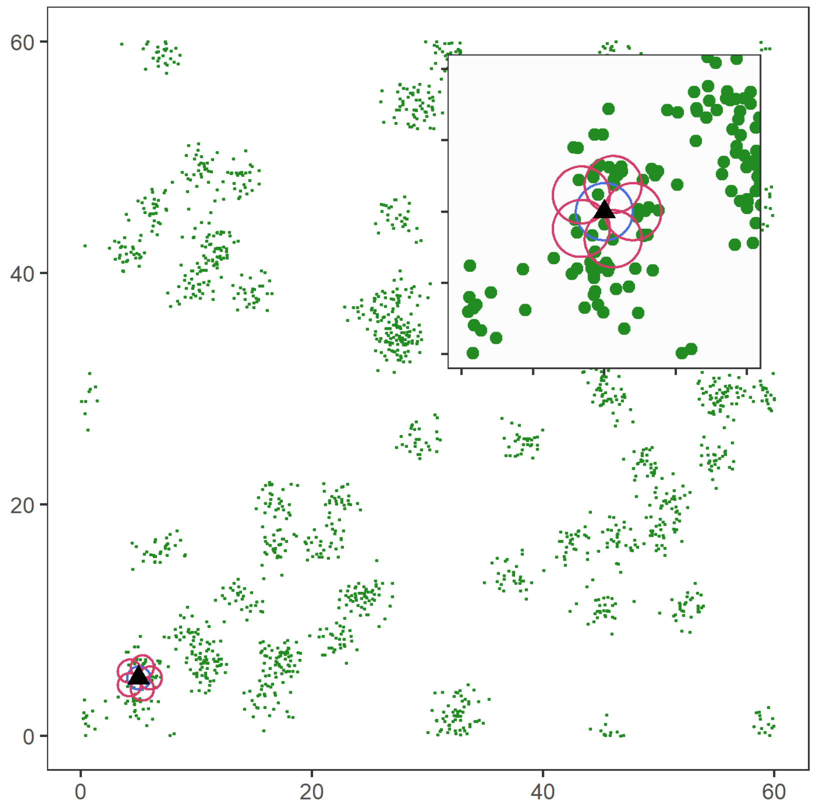

**Figure S2: An example of the resource landscape used in our simulations.** Our simulation's resource landscape consists of 60 randomly distributed clusters of food items ('resource patches'), with 1800 discrete food items divided among the clusters (30 items per cluster). The landscape is a square of 60 units per side, with wrapped boundaries (i.e., a torus). The food item density in our scenarios is 0.5 food items per unit area. Items are distributed around the centre of each cluster, within a standard deviation of 1.0 unit. Items, once consumed by foragers, are unavailable for a fixed number of timesteps (the regeneration time $R$, expressed in terms of the foragers' generation time), after which they regenerate in the same location. While regenerating (i.e., unavailable). While regenerating, items cannot be sensed by foragers. The sensory ranges of individuals ($d_S$) are shown for each potential step (**red circles**, including the current location: **blue circle**). Food item clustering means that available items, as well as foragers handling a food item (handlers) are good indicators of the location of a resource cluster.

### 2 Comparing ecological outcomes across parameter combinations

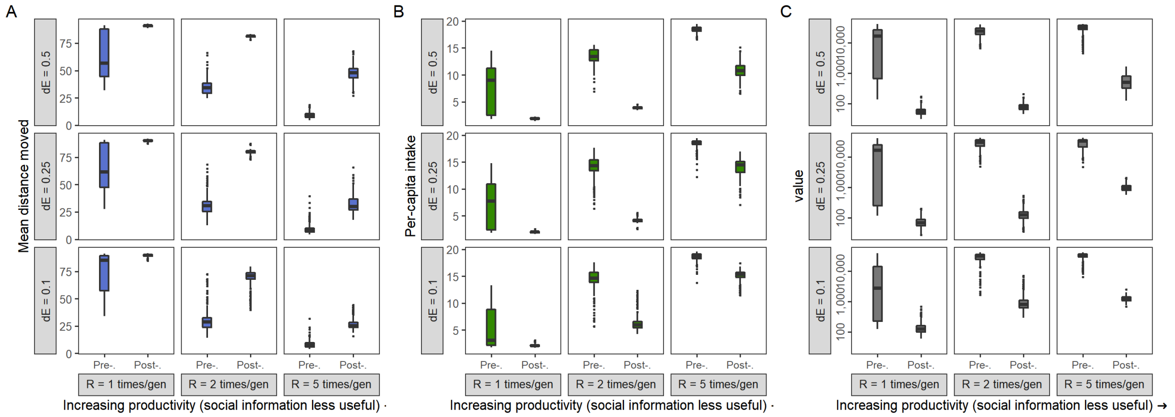

**Figure S3: Rapid changes in ecological outcomes following pathogen introduction.** The introduction of the infectious pathogen leads to rapid evolutionary changes in movement strategies (see Figures 1 and 4; main text) across most combinations of landscape productivity and infection cost. In all combinations where there is rapid evolutionary shift in social-movement strategies, there is a similar change in the population's ecological outcomes: more movement, less intake, and fewer associations. Each subplot in each panel shows the mean and standard error of the per-capita values for **(A)** distance moved, **(B)** intake, **(C)** number of associations, or encounters, with other individuals. In a number of ecological parameter combinations, the evolutionary shift in movement strategies due to pathogen introduction causes reductions in intake (panel B) that are comparable to a halving of landscape productivity. Means and standard deviations are shown before (G = 3,000) and after (G = 3,500) pathogen introduction; each data point represents 10 replicates of the relevant parameter combination.

### 3 The effect of modelling choices

Modelling choices can have a substantial effect on the outcomes of simulations with multiple, complex interactions among components (Scherer et al. 2020, Gupte et al. 2021, Netz et al. 2021). We show the effect of varying implementation on two key aspects of our model: (1) where individuals are initialised, or 'born', on the landscape (natal dispersal), (2) how the infectious pathogen imposes fitness costs.

#### 3.1 Global natal dispersal of individuals

Some models initialise the individuals in each new generation at random locations on the landscape (see e.g. Gupte et al. 2021); this can be called 'global' natal dispersal. This is a reasonable choice when modelling animals during a specific stage of their life cycle, such as after arriving on a wintering or breeding site after migration. Our default choice, on the other hand, is 'local' natal dispersal, where individuals are initialised close to their parent's last position. This is also defensible, as many organisms do not disperse very far from their ancestors. When animals do not disperse very far, they may not evolve movement rules that can be generalised across all landscape conditions, especially when the landscape is ecologically heterogeneous. Instead, animals may adapt their strategies to the local conditions which they inherit from their parents ("ecological inheritance": Badyaev and Uller 2009).

Successful individuals are likely to have more offspring than unsuccessful individuals, and successful individuals are likely to be found — in our simulation and in real natural systems — on or near profitable resource patches. This means that many individuals are initialised near profitable patches. In this case, and because of the sparse distribution of resource patches on the landscape, individuals adapt to tolerate their many neighbours (who are often kin), as avoiding them would lead to also moving away from a profitable patch.

By forcing animals in each new generation to encounter ecological circumstances potentially different from those of their parents, implementing global dispersal can help investigate whether animals' evolved movement strategies are truly 'optimal' at the global scale (Gupte et al. 2021). We implementated global dispersal by running 10 replicates of each parameter combination (9 combinations of $\delta E$ and \$R; 90 simulations in all), with dispersal set to 10. This means that individuals' initial positions are drawn from a normal distribution with standard deviation = 10, centred on the location of their parent (see Figure 4; blue circles).

**Evolutionary outcomes of the global dispersal implementation**

In the global dispersal scenario (see Figure 5), there is a marked difference in which social movement strategy is evolved before pathogen introduction. Since individuals are initialised relatively far away from their parent's position, they encounter potentially very different ecological conditions, both in terms of the number of other individuals, and the local availability of food items.

As a result, most individuals evolve a 'handler tracking' social movement strategy before the introduction of the novel pathogen. This strategy allows individuals to gain the benefits of social information on the location of a resource patch (of which handlers are an indirect cue), while avoiding potential competitors, as well as potentially moving away from areas without many food items.

After pathogen introduction, there is a rapid evolutionary shift in social movement strategies, similar to the shift seen in our default implementation of local dispersal. However, these shifts only occur under conditions where the cost of infection is apparently greater than the value of using social information to find food items. In brief,

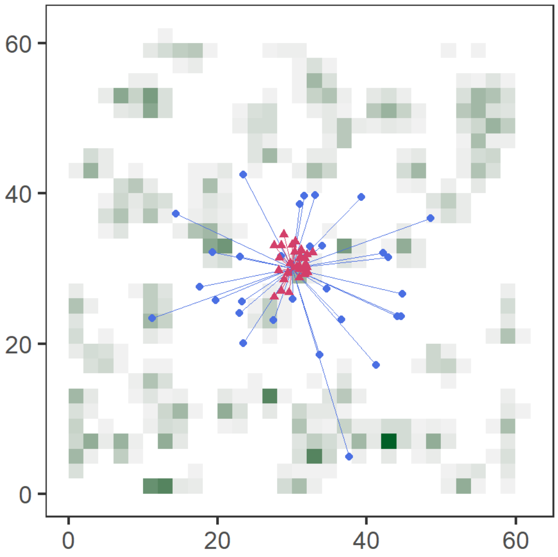

**Figure S4: Differences between local and global dispersal.** Initialising individuals in each new generation within a standard deviation of 10 units around their parent (**blue**; parent at [30, 30]) places can lead them to encounter potentially very different ecological, and social, circumstances from those of their parent. In contrast, individuals initialised close to their parents (within a standard deviation of 2 units; **red**) encounter very similar conditions as their parent. The latter also leads to substantial competition among kin. We used 10 units to represent (nearly) global dispersal, and 2 units to represent local dispersal; this is controlled by the simulation parameter *dispersal*, which takes a numeric argument.

*(1)* when the benefits of social information cannot compensate for the costs of infection risk ($\delta E$ = 0.5; $\delta E$ = 0.25, and $R$ = 1, 2), the agent avoiding strategy becomes more prevalent, similar to the local dispersal case. *(2)* When the costs of infection are lower than the benefits of social information, or when the resource landscape's productivity can offset the cost of infection, the handler tracking strategy persists as the dominant strategy (see Figure 4).

#### Ecological consequences in the global dispersal implementation

In the global dispersal implementation, there is little to no change in population-level ecological outcomes — mean distance moved, mean per-capita intake, and the mean number of associations — following pathogen introduction. This is despite a shift in evolved social movement strategies under some combinations of ecological parameters (i.e., productivity and infection cost). This is likely because a large part of individual's lifetimes (at low $R$, up to 90 timesteps), are spent moving, likely as they search for resource clusters. Since intake depends on finding these clusters, and associations mostly take place at or near resource clusters, these are also reduced compared to our local dispersal implementation.

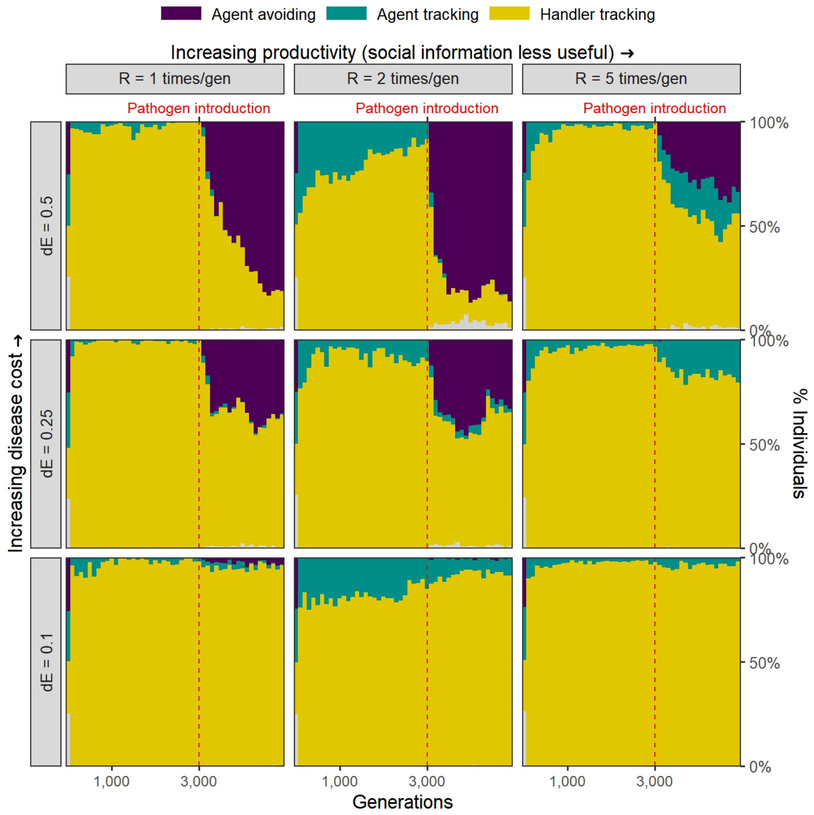

**Figure S5: Pathogen introduction triggers similar evolutionary changes under global dispersal as under local dispersal.** In our alternative, global natal dispersal implementation, the handler tracking strategy is the dominant strategy across most parameter combinations prior to pathogen introduction. Following pathogen introduction, there is a rapid shift in the mix of movement strategies under some ecological conditions. When the cost of infection is greater than the apparent benefit of social information, the agent avoiding strategy becomes more common. When infection costs are low ($\delta E = 0.1$), pathogen introduction does not alter the mix of movement strategies, and the handler tracking strategy continues to be the most common strategy.

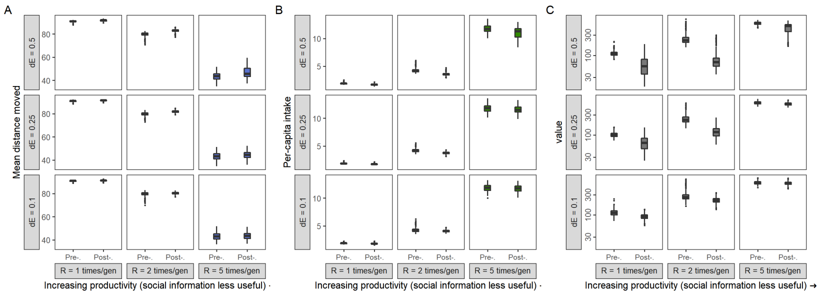

**Figure S6: Little to no change in ecological outcomes when implementing global dispersal.** Despite strong and rapid evolutionary shifts in social movement strategies, the ecological outcomes for populations with global natal dispersal are very similar before and after the introduction of the infectious pathogen. Each subplot in each panel shows the mean and standard error of the per-capita values for **(A)** distance moved, **(B)** intake, **(C)** number of associations, or encounters, with other individuals. Means and standard deviations are shown before (G = 3,000) and after (G = 3,500) pathogen introduction; each data point represents 10 replicates of the relevant parameter combination.

#### 3.2 Infection cost as a percentage of intake

**Default implementation: Infection costs and intake are independent**

In our model's default implementation, the infectious pathogen imposes a direct cost, $\delta E$, on individuals, in each timestep that they are infected. For an individual with intake $N$, the net energetic gain $E$ after being infected by a pathogen for $t$ timesteps is $E = N - (\delta E \times t)$.

In this scenario, *infection costs are independent of intake*.

**Alternative implementation: Infection costs as a percentage of intake**

In an alternative implementation, the infectious pathogen may be considered to reduce an animal's ability to process intake, or to require a portion of daily intake to resist. Such an implementation is used in . . .

For an individual with intake $N$, the net energetic gain $E$ after being infected by a pathogen for $t$ timesteps is $E = N \times (1 - \delta E)^t$.

**Comparing cost structures across implementations**

Naturally, the two cost structures are not easy to compare.

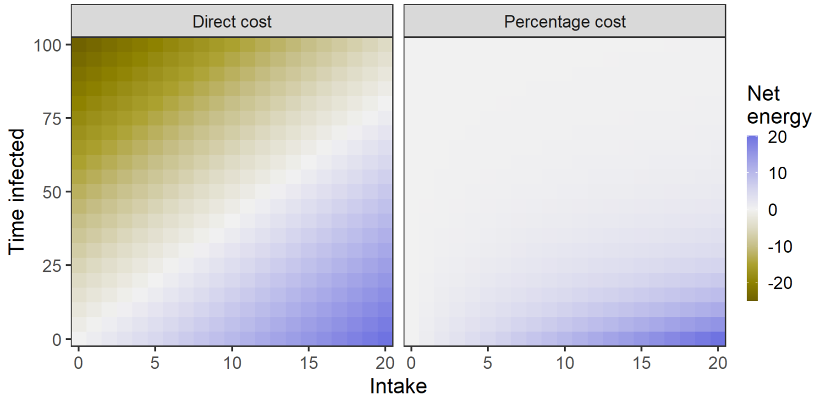

**Figure S7:** Calculated net energy for different combinations of intake and time infected. In the *Direct cost* scenario, and with a $\delta E$ of 0.25 (shown here), which is our default implementation, an individual foraging on an item (handling time = 5 timesteps) would gain 1.0 unit of intake, and lose 1.25 units of energy in that same period if it were infected, for a net energy balance in that period of -0.25. Individuals' energetic balance is normalised (0 – 1) with reference to the lowest value in each generation. Here, individuals' infection cost is *independent* of their intake. In the *Percentage cost* scenario, individuals' infection costs are linked to their intake. For a per-timestep 5% loss of intake (shown here), individuals infected for >25 timesteps already have a net energy balance close to, but never less than, zero. In this implementation, individuals' energy balances are *not normalised* with reference to the lowest net energy, as no individual's energy is ever less than zero.

#### Evolutionary outcomes of the percentage cost implementation

The social movement strategies evolved prior to pathogen introduction are identical to those seen in our default implementation. This is because the percentage cost implementation differs from the default only after the pathogen is introduced.

After pathogen introduction, there is a rapid evolutionary shift in movement strategies. This shift is similar to that in our default implementation, but the strategies evolved are different. The handler tracking strategy becomes common across parameter combinations. However, when the costs of infection are relatively high (7.5%), and the usefulness of social information is limited by the abundance of food items (R = 5), the agent avoiding strategy forms about one fourth of the population mixture of social movement strategies.

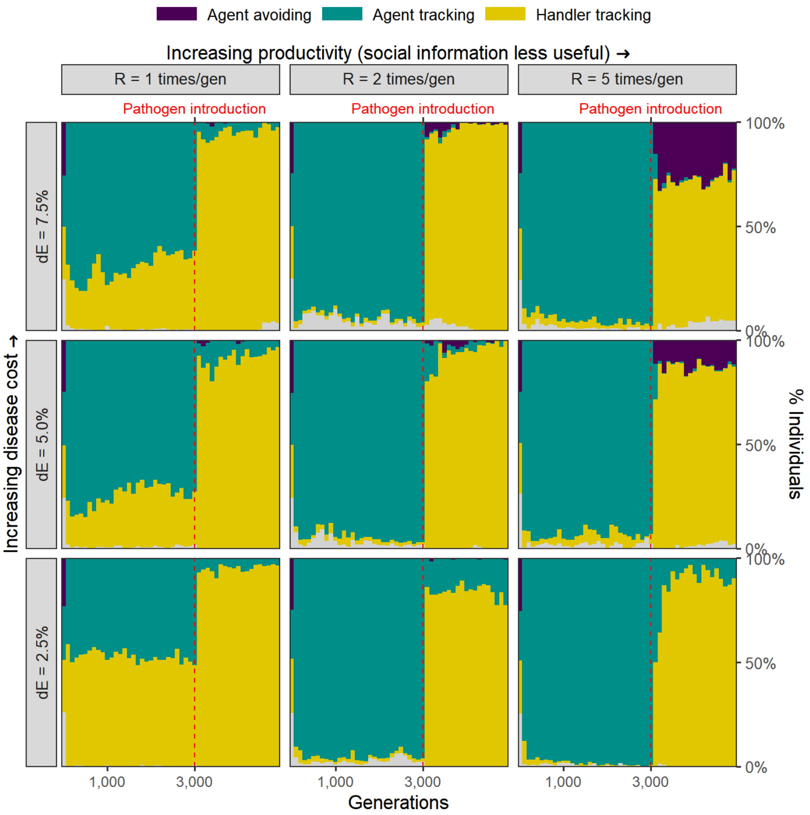

**Figure S8: Rapid evolutionary change, but different evolutionary outcomes, in an alternative implementation of disease costs.** In our alternative, percentage costs implementation of the infectious pathogen, there is a rapid shift in the mix of movement strategies after pathogen introduction. The handler tracking strategy becomes common across all parameter combinations. Only when the costs of infection are relatively high (7.5%), and the usefulness of social information is limited by the abundance of food items (R = 5), does the agent avoiding strategy form about one fourth of the population mixture of social movement strategies.

#### Ecological consequences in the percentage cost implementation

Surprisingly, the implementation of a different cost structure for the novel, infectious pathogen does not affect ecological, population level outcomes when compared with outcomes in our default implementation of direct costs. Across parameter combinations where there is a rapid evolutionary transition from agent tracking to handler tracking as the dominant strategy, there is also an increase in distance moved, a reduction in intake, and a reduction in associations. Notably, the reductions in per-capita intake following pathogen introduction are similar to a halving of landscape productivity (as in the default implementation), and there is a comparable drop in the number of pairwise associations among individuals.

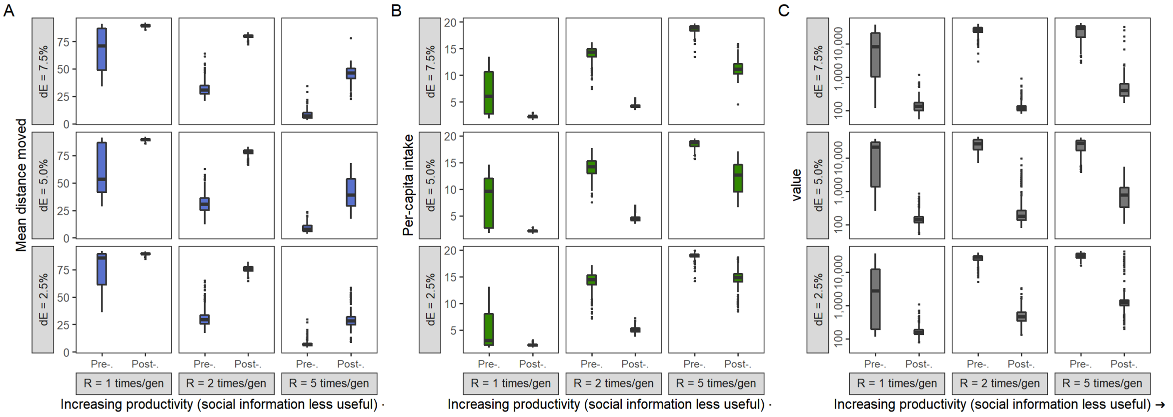

**Figure S9: Rapid ecological changes accompany evolutionary shifts in an alternative implementation of disease costs, and are similar to the default implementation.** In the alternative percentage-costs implementation of the infectious pathogen, the outcomes are very similar to those in our default implementation of direct costs. Across most paramter combinations, there is an increase in movement, a reduction in intake, and a reduction in associations with other foragers. Each subplot in each panel shows the mean and standard error of the per-capita values for **(A)** distance moved, **(B)** intake, **(C)** number of associations, or encounters, with other individuals. Means and standard deviations are shown before (G = 3,000) and after (G = 3,500) pathogen introduction; each data point represents 10 replicates of the relevant parameter combination.

#### 3.3 Sporadic introduction of infectious pathogens

We implemented a variant of our main model, in which the infectious pathogen is introduced only sporadically after the first introduction event (at G = 3,000). Specifically, we modelled probabilistic introduction of the pathogen in each generation following the initial introduction. We call the per-generation probability of a novel pathogen introduction event the 'spillover rate'. We ran 10 replicates each of this model variant and examined whether there was a similar evolutionary shift in social movement strategies as seen in our default implementation. Since it is the main parameter of interest, we ran this model variant for three values of the spillover rate: 0.05, 0.1, and 0.25. Instead of examining the joint effect of landscape productivity and cost of infection as well, we only examined the effect of infection cost, implementing three different variants with an infection cost $\delta E$ of 0.1, 0.25, and 0.5. We kept all other model parameters similar to the default scenario of our main model, and importantly, considered only a landscape productivity $R$ of 2. Cross-species novel pathogen introductions are likely to become more common with climate change, and so we chose these spillover rate values to represent different scenarios under altered global regimes of pathogen transfer. Our model's default implementation may be seen as an extreme case of the models considered here, with a spillover rate of 1.0.

##### Model implementation

In our model code, the sporadic introduction is implemented by drawing the number of generations until the next pathogen introduction event from a geometric distribution whose probability parameter is given by the spillover rates described above. Zero values are handled by converting them into ones. At our lowest spillover rate, up to 100 generations could pass between pathogen introductions, while at our highest rates, there are rarely more than 10 generations between introductions.

##### Evolutionary outcomes of the percentage cost implementation

The social movement strategies evolved prior to pathogen introduction are identical to those seen in our default implementation, as expected. However, following pathogen introduction, we found that there was little to change in the population-level mixture of movement strategies in this model variant (see figure). This is regardless of the probability of a novel pathogen introduction (our so-called 'spillover rate'), and the cost of infection by a pathogen. Across the simulation, the commonest social movement strategy remains 'agent tracking', i.e., preferring locations with multiple individuals regardless of their foraging status.

Since there is little to no change in social movement strategies, we did not expect nor find changes in ecological outcomes.

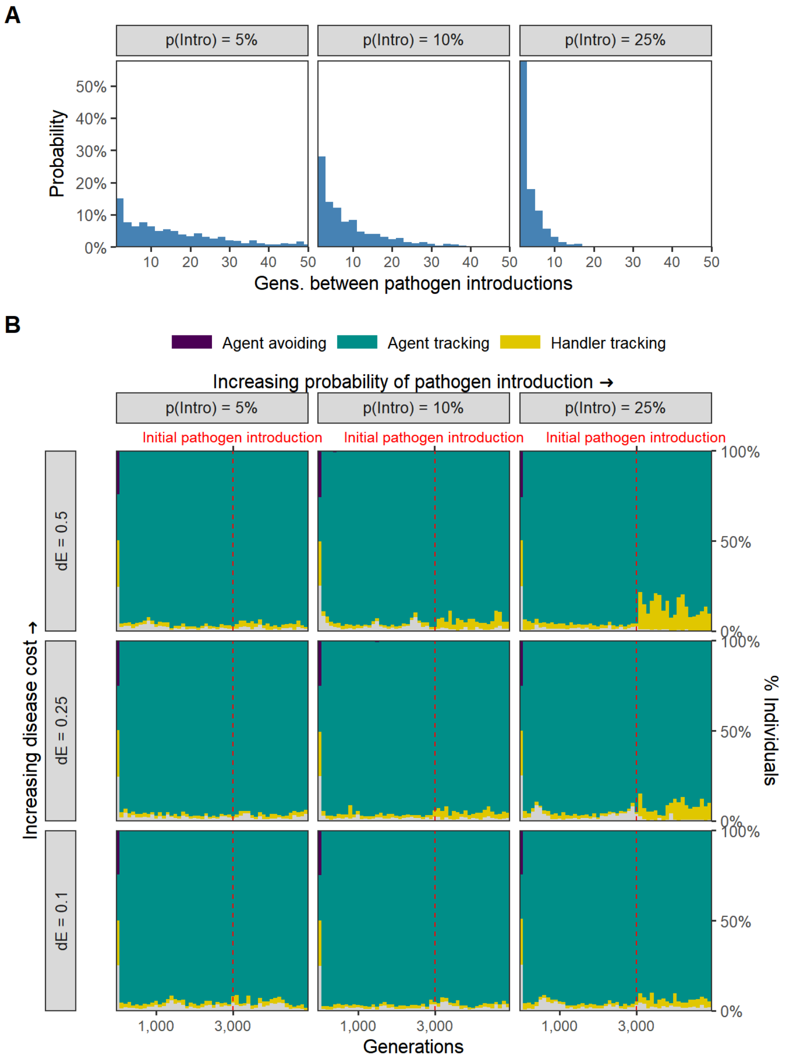

**Figure S10: No evolutionary change in social movement strategies when novel pathogen introduction events are relatively uncommon. (A)** In our alternative implementation of the model, the pathogen is only introduced sporadically after the initial introduction (G = 3,000; red line in panel B). **(B)** When the introductions are relatively rare and sporadic, there is no shift in the mixture of movement strategies after pathogen introduction. The agent tracking strategy remains common across parameter combinations.

Scherer, C. et al. 2020. Moving infections: Individual movement decisions drive disease persistence in spatially structured landscapes. - Oikos 129: 651–667.
